# Neocortical Feedback Biases Hippocampal Reactivation Toward Unconsolidated Memories in a Predictive-Coding Model of Systems Consolidation

**DOI:** 10.64898/2026.09.29.755342

**Authors:** Saighi Paul

## Abstract

During sleep and quiet wakefulness, the brain spontaneously revisits neural activity patterns associated with previous experience, a phenomenon known as reactivation, or replay. Reactivation is thought to support systems consolidation, the process by which memory retrieval becomes progressively more dependent on neocortical circuits and less dependent on the hippocampus. Importantly, reactivation is not indiscriminate: some memory traces, such as newly acquired or weakly learned ones, are preferentially reactivated.

These observations have motivated the development of computational models accounting for the spontaneous reactivation of memory traces, their prioritisation for replay, and their transfer between the hippocampus and neocortex. In parallel, predictive-coding (PC) models have shown how computations attributed to hippocampal and neocortical circuits can arise from local prediction-error dynamics. Here, we connect these two lines of work by showing how autonomous intrinsic dynamics in coupled PC networks can transfer stored memory content between neural systems.

The hippocampus and neocortex are modelled as two coupled PC associative memory networks, with asymmetric coupling motivated by reported interactions between hippocampal area CA1 and the prefrontal cortex. This coupling establishes a closed-loop hippocampal-neocortical dialogue in which hippocampal activity drives neocortical learning, while neocortical feedback biases subsequent hippocampal activity toward content for which neocortical support remains weak. We prove that this feedback steers hippocampal activity toward states of higher neocortical free energy, so that prioritisation emerges from the dialogue itself rather than from an explicit scheduler.

In our model, neocortical learning relies exclusively on spontaneous hippocampal activity, without access to the initial training corpus. Nevertheless, we show that the neocortical network acquires correlated memory content faster than direct training on randomly sampled memory patterns. This advantage appeared for sufficiently correlated patterns, grew with their correlation, and persisted when new patterns were incrementally added during transfer.

Together, these results provide a PC account in which neocortical feedback organises hippocampal reactivation to support efficient transfer of memory content during systems consolidation.

## 1 Introduction

A large fraction of the brain’s metabolic budget is devoted to intrinsic activity, that is, ongoing neuronal dynamics not directly driven by current sensory input or behavioural demands [1, 2]. During sleep or quiet rest, intrinsic activity can reinstate activity patterns associated with prior waking experience, a phenomenon known as neural reactivation [3–5]. These reactivation events can be coordinated across hippocampal and neocortical circuits, forming part of the hippocampal-neocortical dialogue [6–8]. The term replay is also used to designate neural reactivation, but can refer more narrowly to the reactivation of ordered temporal sequences of experience-related activity patterns. Reactivation is therefore used throughout this study as the broader term.

Classical accounts propose that the hippocampal-neocortical dialogue supports systems consolidation by gradually shifting the neural basis of memory retrieval from hippocampal to neocortical circuits [7, 9–11]. Systems consolidation is therefore often described as a redistribution of memory representations across brain systems, or as a form of memory transfer whereby the neocortex progressively acquires memory content initially encoded in the hippocampus [10, 12, 13].

A substantial literature indicates that the reactivation events underlying systems consolidation preferentially target certain memory traces over others [5,14]. Among other factors, traces that are recent, that correspond to novel experiences, or that were weakly learned are prioritised for reactivation [3, 5, 14, 15]. The precise neural mechanisms inducing this prioritisation remain poorly understood [5, 14].

Reactivation and systems consolidation have motivated a broad range of computational models, from complementary learning systems [13] to models of offline or autonomously generated reactivation [16–18], generative replay [19, 20], and models in which reactivation or consolidation is prioritised according to utility, context, or expected benefits for generalisation [21–25]. These accounts differ in what drives reactivation or consolidation, such as memory recency, attractor competition, context, reward, or expected utility, and in what information is transferred.

In this work, we focus on a complementary account based on a single candidate mechanism, grounded in a specific communication motif between the hippocampus and neocortex. The communication motif we consider has been characterised between the prefrontal cortex (PFC) and hippocampal area CA1, and combines two directional influences: recruitment of PFC activity by CA1 and suppression of CA1 activity associated with independent PFC reactivation. During sleep, CA1 activity is followed by increased firing in functionally coupled PFC neurons [26]. Conversely, when PFC populations reactivate independently, without coincident CA1 reactivation, CA1 neurons that normally participate in coordinated CA1–PFC events are preferentially suppressed [27].

We interpret these findings as indicative of an asymmetric and selective communication scheme between the systems: hippocampal reactivation recruits specific prefrontal populations, whereas prefrontal reactivation suppresses the corresponding hippocampal representation.

Wang, Poe, and Zochowski proposed a functional interpretation of such a motif, in which neocortical inhibitory feedback regulates hippocampal reactivation according to the degree of transfer [28]. In their network model, hippocampal reactivation drives cortical activity, and slow cortical plasticity progressively builds a corresponding cortical representation. The cortex also projects diffusely onto hippocampal inhibitory interneurons, but this feedback becomes effective only once the cortical representation has matured. Reactivation of a consolidated trace then evokes strong, self-sustained cortical activity that transiently suppresses hippocampal activity. Reactivation of consolidated traces is thus shortened or abolished, whereas traces that still lack cortical support continue to reactivate. As transfer proceeds, reactivation becomes increasingly biased toward content that has not yet been transferred.

Here, we extend this account in three respects. First, we formulate the interacting hippocampal and neocortical memory systems within the predictive-coding (PC) framework. Second, we replace diffuse feedback inhibition with selective inhibition, consistent with the experimental observations mentioned above. Third, we demonstrate the functional role of this selective inhibition in organising memory transfer, showing that the resulting dynamics enable fast transfer of correlated memory patterns.

In the proposed model, neocortical feedback inhibition biases reactivation away from hippocampal content already supported cortically and toward content that remains poorly transferred. This dynamic directs reactivation toward the state in the hippocampal memory subspace that maximises settled neocortical free energy. Neocortical plasticity then reduces the corresponding free energy. The resulting architecture therefore gives rise to prioritised reactivation and transfers memory content as consolidation proceeds.

As a first step toward a more biologically detailed model, we study a simplified setting in which the hippocampus and neocortex are each represented by a single linear associative memory network. The neurons are rate units rather than spiking neurons, simplifying the analysis.

Both networks are further assumed to share the same neural code and to communicate through a one-to-one pathway, in which each hippocampal neuron is connected to a single corresponding neocortical neuron. In the brain, by contrast, hippocampal-neocortical connectivity is not one-to-one, and the two systems interact through polysynaptic routes as well as direct projections [29–32]. Moreover, hippocampal indexing theory proposes that the hippocampus stores an index of neocortical activity patterns rather than a copy of them [33, 34]. We motivate these simplifications in the Discussion.

## 2 Model

The model consists of two coupled associative-memory predictive-coding networks (PCNs): a teacher network, corresponding to the hippocampus, and a student network, corresponding to the neocortex. In isolation, each network performs inference: its neuronal activity evolves to minimise the network’s own free energy. In a trained network, this allows a stored pattern to be retrieved from a partial cue. Learning dynamics are derived from the same free energy minimisation principle, but through changes in synaptic weights. The full architecture adds an asymmetric coupling between the teacher and student networks.

Memory storage occurs in two successive phases. The teacher first stores externally acquired activity patterns, corresponding to memory encoding during awake activity. Autonomous intrinsic activity of the coupled system then transfers the teacher’s memory content to the student network.

### 2.1 A single predictive-coding associative memory

Tang et al. introduced the implicit covariance-learning PCN (implicit covPCN), which constitutes the basic building block of our architecture [35]. The implicit covPCN was developed as a more biologically plausible alternative to the explicit covPCN introduced in earlier works [36–38]. Here, we present the linear version of the implicit covPCN.

Both the teacher *T* and the student *S* are implicit covPCNs with dense recurrent connectivity and no self-connections (autapses).

We use *α* ∈ {*S, T*} to index either network. For activity *x*_*α*_ ∈ ℝ^*d*^ and recurrent weights *W*_*α*_, we define the prediction-error operator and recurrent error

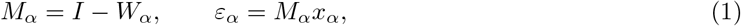

and the network free energy,

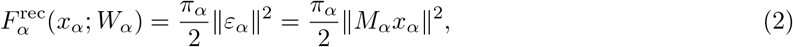

where *π*_*α*_ *>* 0 is the fixed precision assigned to recurrent prediction errors. Following Tang et al., we interpret the precision-weighted prediction-error objective as a simplified variational free-energy functional under Gaussian assumptions. We refer to it below simply as free energy.

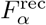 accounts only for the network’s internal prediction error generated by its recurrent connectivity. Two network dynamics operating on distinct timescales reduce 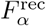: fast *inference*, through changes in neuronal activity, and slow *plasticity*, through changes in synaptic weights,

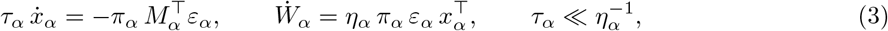

with activity time constant *τ*_*α*_ and learning rate *η*_*α*_. Weight updates and gradients with respect to the weights are understood to act only on the off-diagonal entries; self-connections remain frozen at zero.

The prediction may optionally pass through a pointwise nonlinearity *f*, giving *ε*_*α*_ = *x*_*α*_ − *W*_*α*_*f* (*x*_*α*_) [35]. We take the linear case *f* = id throughout this study. In addition, as introduced in the next section, the teacher activity is normalised such that ∥*x*_*T*_∥ = 1 after each integration step.

Each network is trained according to the plasticity dynamics in Eq. (3), either during interleaved clamping or through intrinsic activity during memory transfer (Section 2.3).

In a fully trained network *α*, the stored content becomes the kernel of its error operator,

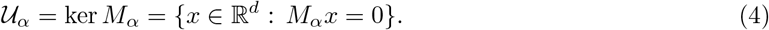

*U*_*α*_ can therefore be interpreted as the network’s memory subspace. Every stored pattern *m*^(*p*)^ satisfies *M*_*α*_*m*^(*p*)^ = 0 and has zero recurrent free energy 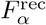 [35].

Because *U*_*α*_ is a linear subspace, any linear combination of stored patterns is also a zero-energy fixed point. The network therefore stores a flat memory manifold rather than the discrete point attractors of classical Hopfield networks [39, 40].

### 2.2 Coupling the teacher and the student

Fig. 1 shows the full architecture formed by coupling the student *S* and teacher *T*.

**Fig 1.**
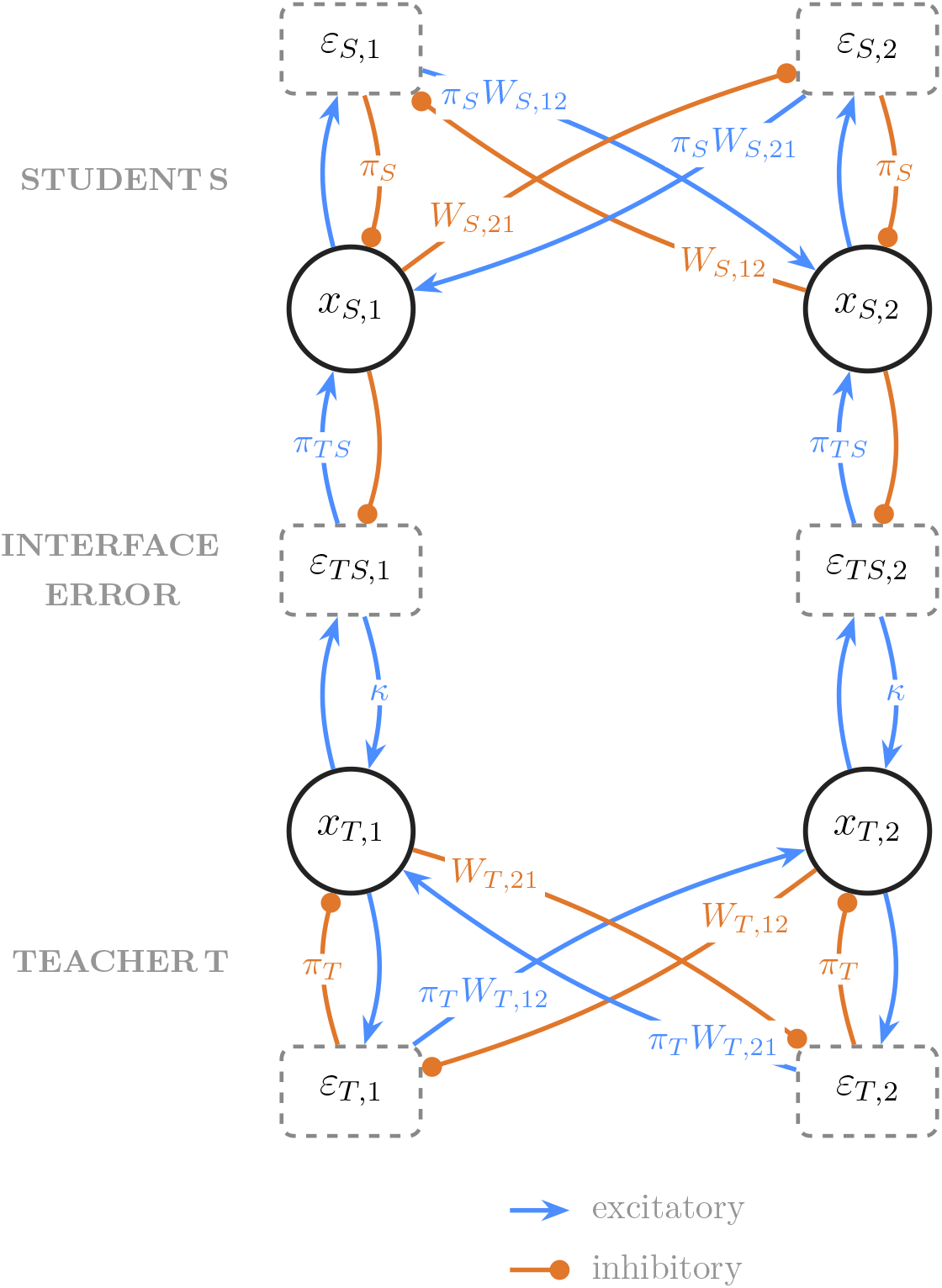
Model architecture (two neurons per population). Student *S* (top) and teacher *T* (bottom): value neurons *x*, recurrent errors *ε*_*S*_, *ε*_*T*_, and the one-to-one interface error *ε*_*T S*_ = *x*_*T*_− *x*_*S*_. Connections without labels have magnitude 1. *κ* denotes the scalar strength of an excitatory synapse, whereas in classical PC networks, the top-down influence from the interface error is inhibitory.

The two networks are coupled through an *interface-error* population, with each error unit receiving input from exactly one teacher neuron and its corresponding student neuron:

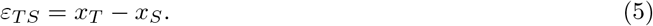

Teacher and student activities are therefore expressed in a common coordinate system: the *i*-th teacher neuron and the *i*-th student neuron encode the same component of a memory pattern. We return to this architectural choice in the Discussion.

Without the interface influence, the activities of both the teacher *T* and the student *S* follow the inference dynamics in Eq. (3) and descend their respective recurrent free energies 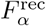.

For the teacher, the coupling adds a direct interface-error drive, giving

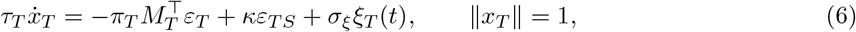

where *κ* is the teacher-side coupling gain. During intrinsic activity, *κ >* 0, so the interface-error population excites the teacher. In conventional PC, minimising the interface error with respect to *x*_*T*_ would instead contribute −*π*_*T S*_*ε*_*T S*_ to the teacher dynamics. The teacher-side pathway therefore reverses the conventional sign. The term *σ*_*ξ*_*ξ*_*T*_ (*t*) provides low-amplitude stochastic input, with *ξ*_*T*_ (*t*) denoting standard Gaussian white noise and *σ*_*ξ*_ its amplitude.

After each integration step, teacher activity is normalised to unit norm to prevent unbounded activity. This operation idealises fast population gain control. In biological circuits, divisive normalisation can constrain response magnitude through several mechanisms, including feedforward and recurrent inhibition [41, 42].

The student side retains the conventional PC dynamics. Student activity minimises its recurrent free energy and receives an excitatory influence from the interface-error population,

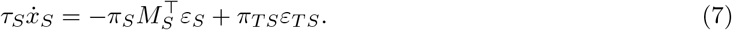

Both contributions to Eq. (7) are gradient-descent terms on the student free energy, which can therefore be expressed as

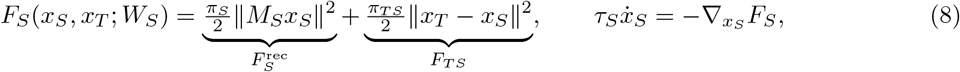

where *π*_*T S*_ *>* 0 is the precision of the interface prediction error. Activity and plasticity evolve concurrently in the simulations.

The overall influence between the student *S* and the teacher *T* requires careful interpretation. Although *κ >* 0 denotes an excitatory connection from the interface-error population to the teacher, the student enters *ε*_*T S*_ = *x*_*T*_ − *x*_*S*_ with a negative sign, inhibiting the interface error. The combined pathway *x*_*S*_ → *ε*_*T*_ → _*S*_ *x*_*T*_ is thus inhibitory: increasing student activity reduces the excitatory error drive reaching the teacher. This effective top-down inhibition is the model counterpart of prefrontal suppression of hippocampal activity. In a conventional PC network, the interface error would instead inhibit the lower population, producing top-down excitation through disinhibition.

### 2.3 Model dynamics

Our transfer protocol consists of two phases. As shown in Fig. 2, the first establishes the teacher’s memory subspace, and the second transfers its content to the student without re-presenting the original data.

**Fig 2.**
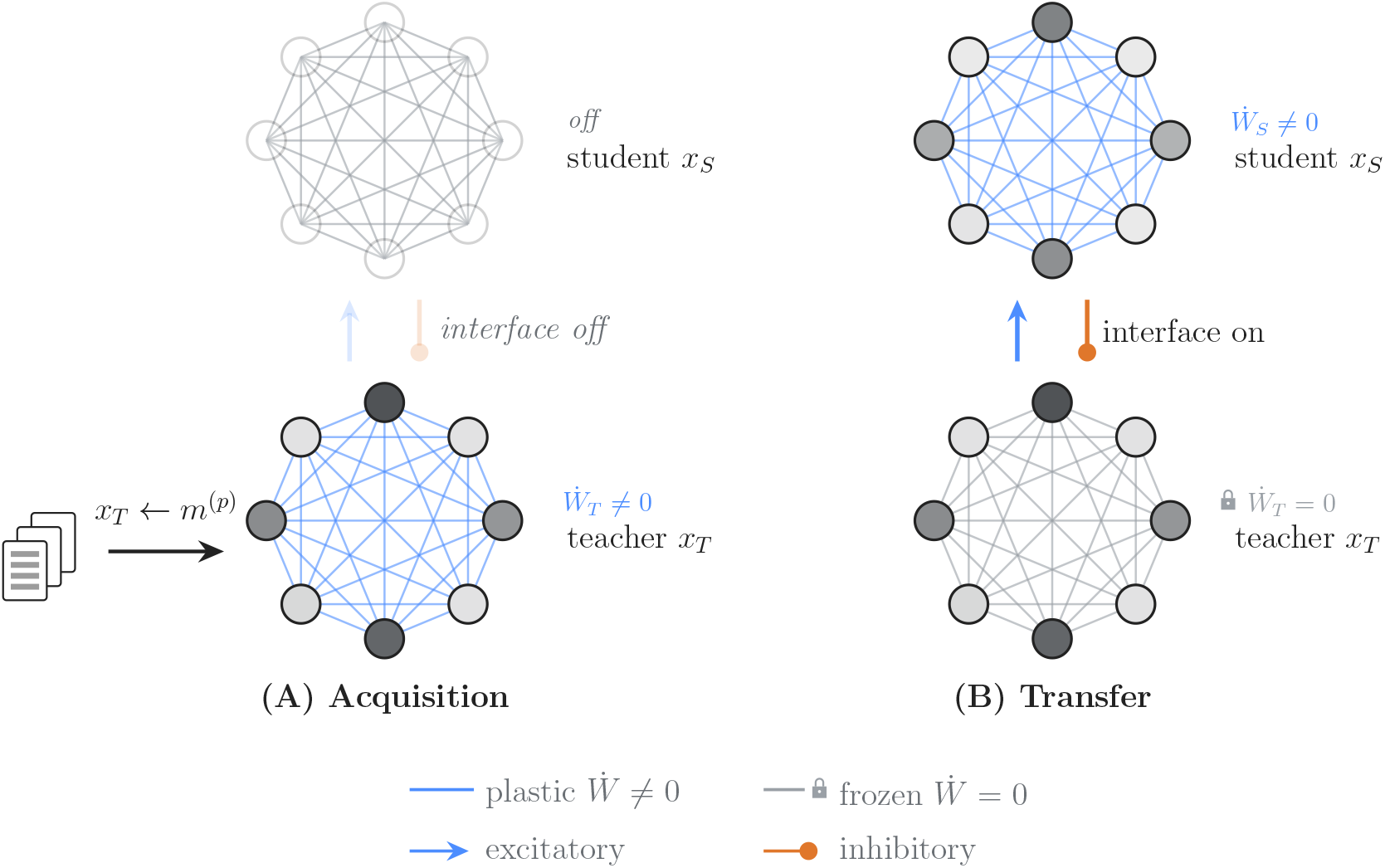
Initial memory acquisition and transfer dynamics. **A**, The teacher is trained on the memory corpus via interleaved training. **B**, Subsequent intrinsic dynamics transfer memory content from the teacher to the student.

During initial storage (Fig. 2A), teacher activity *x*_*T*_ is clamped to memory patterns *m*^(*p*)^, presented in interleaved order, while *W*_*T*_ follows Eq. (3). The student and the interface remain inactive. These synaptic updates reduce the teacher’s recurrent prediction error on the presented patterns and establish its memory subspace *U*_*T*_ = ker *M*_*T*_ (Materials and methods, Section 5.2). Interleaved presentation allows the teacher to store correlated patterns while limiting interference between them [13, 43]. Biological hippocampal circuits, by contrast, can rapidly encode individual experiences without the repeated interleaved presentations used here [13, 44]. This ability is thought to depend in part on pattern separation in the dentate gyrus, where granule-cell populations transform correlated inputs into more orthogonal activity patterns [45, 46]. The model lacks this pattern-separation mechanism, so interleaved training is used instead.

During transfer (Fig. 2B), the external clamp on the teacher is removed and the interface is activated. The teacher and student activities follow Eqs. (6) and (7), while *W*_*T*_ remains frozen and only *W*_*S*_ follows the plasticity rule of Eq. (3). The student is never clamped: intrinsic activity induces memory reactivation in the teacher network, which subsequently influences the student through the interface error. The teacher and the student then display similar activity patterns, and recurrent plasticity lets the student learn these reactivated patterns. Implementation details are given in Materials and methods, Section 5.1.

The transfer mechanism assumes that teacher activity remains near the teacher memory subspace. To promote this regime, we impose

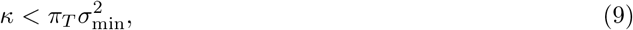

where 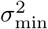 is the smallest nonzero eigenvalue of 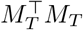. This heuristic promotes, but does not guarantee, confinement of teacher activity near the memory subspace. In contrast, if *κ* is too large, the interface drive can overcome the teacher’s recurrent restoring dynamics, pushing activity into off-memory-subspace directions and impairing transfer.

## 3 Results

### 3.1 Reactivation and memory content transfer

We first examined whether intrinsic activity in our model could reactivate and transfer stored memory content from the teacher network to the student network.

We stored three unit-normalised MNIST patterns [47] in a 784-unit teacher network, froze the teacher’s recurrent weights, and started the coupled dynamics. During this intrinsic phase, neither network received external input. Low-amplitude Gaussian noise was applied to the teacher activity to seed the dynamics. The results are shown in Fig. 3 (Materials and methods, Section 5.4.1).

**Fig 3.**
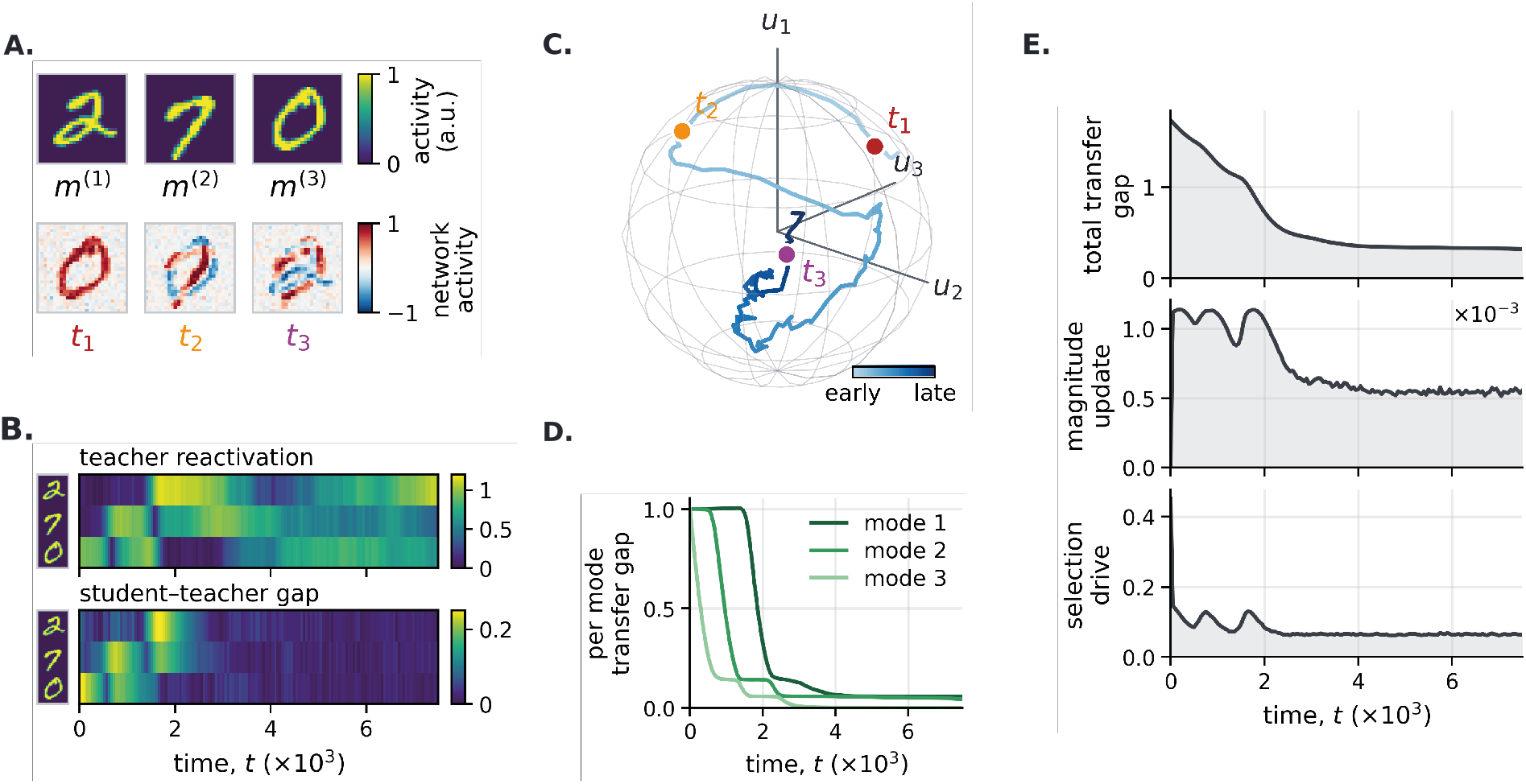
Spontaneous intrinsic activity progressively transfers teacher-supported memory content to the student. The frozen teacher network was coupled to a student network initialised with all recurrent weights set to zero. Low-amplitude Gaussian noise seeded exploration of the teacher memory subspace (Materials and methods, Sections 5.4.1 and 5.4.2). **A**, The three stored MNIST patterns (top; display order 2, 7, and 0) and teacher states at *t*_1_ = 250, *t*_2_ = 1000, and *t*_3_ = 7000 (bottom). **B**, (Top) Memory coefficient magnitude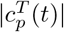, which measures the contribution of each 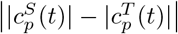; a large gap indicates that the two networks express different activity patterns and a stored pattern *m*^(*p*)^ to teacher activity *x*_*T*_ (*t*). Low values mean that the teacher does not reactivate the memory, whereas large values indicate strong reactivation. (Bottom) Student–teacher magnitude gaps small gap that they display similar activity. **C**, Teacher activity projected onto an orthonormal basis *U*_*T*_ of the three-dimensional teacher memory subspace. The wireframe marks the unit sphere imposed by teacher normalisation, and the coloured markers identify the snapshots in panel A. Basis axes are orthogonal memory-subspace modes, not individual digits. The line colour indicates the temporal order in which the states are explored. **D**, Per-mode transfer gaps, corresponding to the eigenvalues of 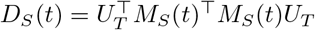. High values identify modes of the teacher memory subspace that are poorly stored by the student, whereas low values indicate modes that are accurately stored. **E**, (Top) Total transfer gap ∥*M*_*S*_*U*_*T*_ ∥_*F*_, which measures how much of the teacher memory subspace remains unsupported by the student. (Middle) Magnitude of student weight updates 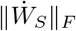. (Bottom) Teacher-side selection drive *κ*∥*ε*_*T S*_∥_2_, the magnitude of the interface-error drive acting on the teacher.

Fig. 3(A, bottom), (B, top), and (C) show that teacher activity is dominated by mixture states rather than by reactivation of individual patterns. This behaviour follows from the linear activation function: the stored patterns span a flat memory manifold containing every signed linear combination of those patterns, including their antipodal states.

Fig. 3(D) and (E) provide direct evidence of memory-content transfer during intrinsic activity. In panel (D), 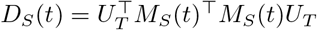 measures the student’s memory-reconstruction error within the teacher memory subspace, which is spanned by the orthonormal columns of *U*_*T*_. Each plotted transfer-gap value is an eigenvalue of *D*_*S*_(*t*). Its associated eigenvector defines an orthogonal mode within the teacher memory subspace, generally a linear combination of several memory patterns, while the eigenvalue gives the squared reconstruction error for that mode (Materials and methods, Section 5.4.2).

The successive reduction of these transfer gaps shows that the student acquires the teacher’s memory subspace one mode at a time, rather than by uniformly reducing the error across the entire subspace. Transfer slows once most of the subspace is stored by the student and remains incomplete over the displayed interval. Longer simulations further reduce the residual transfer gap, although at a much lower rate. At this late stage, the mismatch-driven selection signal is weak and the teacher increasingly relies on noise-induced exploration to sample the remaining poorly represented modes. Despite this incomplete transfer, the isolated-student probes in Fig. 4 show recognisable reconstruction of all three original patterns later during transfer (Materials and methods, Sections 5.4.2 and 5.4.3).

**Fig 4.**
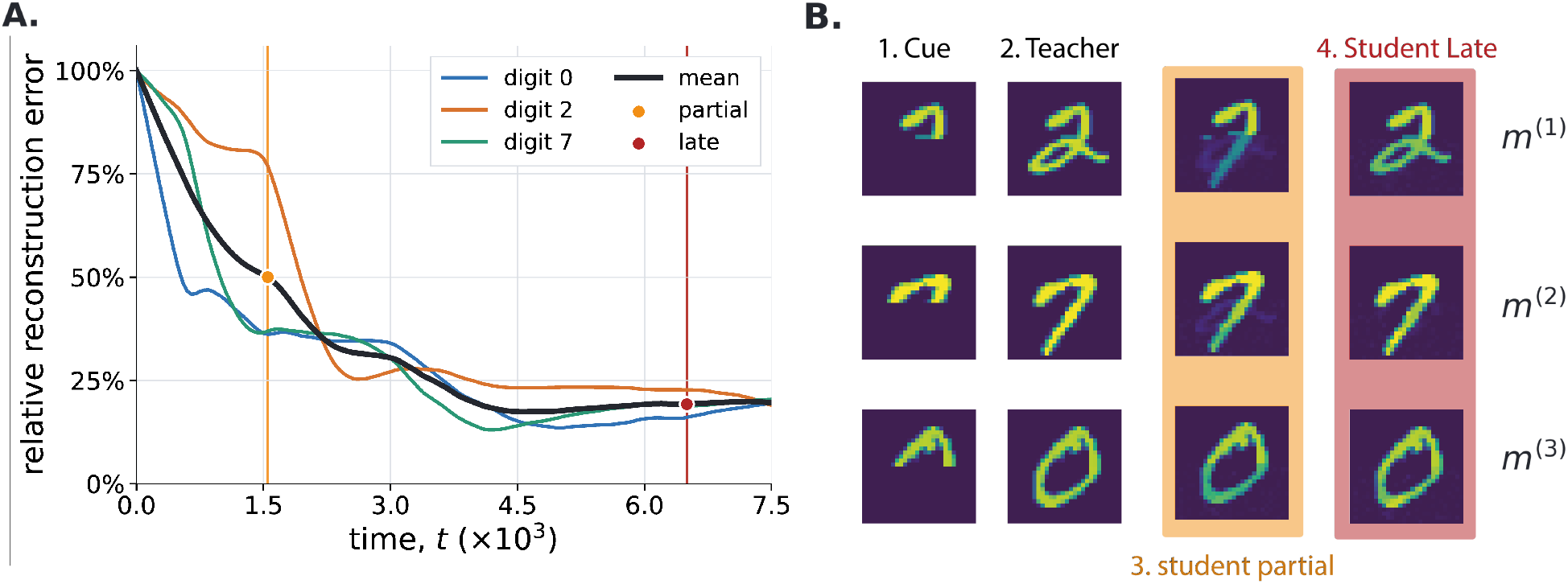
Transferred teacher patterns become reconstructable by the isolated student. The reconstruction residual and clamped-recall protocol are described in Materials and methods, Sections 5.4.2 and 5.4.3, respectively. **A**, Relative recurrent reconstruction error for each stored MNIST pattern during transfer. At each recorded time, the student weights were frozen, its state was set to the complete pattern *m*^(*p*)^, and the relative error *R*_*p*_(*t*) = ∥*m*^(*p*)^ − *W*_*S*_(*t*)*m*^(*p*)^∥_2_*/*∥*m*^(*p*)^∥_2_ was evaluated and expressed as a percentage. Thus, 100% corresponds to the initially empty student with *W*_*S*_ = 0, whereas zero denotes exact recurrent prediction of that pattern. The black curve is the mean over the three digits; orange and red markers identify the partial (*t* = 1550) and late (*t* = 6500) checkpoints used in panel B. **B**, Clamped recall by isolated networks. From left to right, each row shows the partial cue, the teacher reconstruction, and student reconstructions at the partial and late checkpoints. Half of the network neurons were clamped to a given memory pattern, the hidden half was initialised to zero, and the free neurons settled by minimising the network free energy 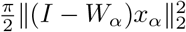, with no teacher–student coupling or ongoing plasticity. Student recall performance is uneven at the partial checkpoint, in agreement with the reconstruction error in panel A. For the late checkpoint, all reconstruction outputs visually approach the teacher completions despite a residual reconstruction error.

Fig. 3(B, bottom) and (E, middle and bottom) show how the signals driving learning and reactivation attenuate during memory transfer. The magnitude of student weight updates decreases as the student– teacher mismatch contracts, revealing a self-limiting learning process: as the student acquires content stored by the teacher, the internal reconstruction error that generates further weight updates is cancelled. In parallel, the teacher-side selection drive, generated by the mismatch-dependent top-down influence of the student on the teacher, also decreases. Three transient peaks accompany the successive reduction of the three eigenvalues in Fig. 3(D), consistent with the sequential acquisition of the three independent modes spanning the rank-three teacher memory subspace. Both signals remain nonzero at the end of the displayed interval due to ongoing noise.

To test whether this mechanism extends beyond transfer from a fixed teacher memory set, we allowed the teacher repertoire to grow during a single transfer run (Fig. 5; Materials and methods, Sections 5.3 and 5.4.4). Each newly added pattern introduced a new high-eigenvalue error mode, while the eigenvalues associated with memory content already transferred to the student remained close to zero. These new transfer gaps then decreased, showing that the same mismatch-dependent dynamics can repeatedly redirect reactivation toward newly unsupported content. This protocol captures a central feature of systems consolidation: the hippocampus rapidly encodes new experiences, while their integration into neocortical circuits unfolds over subsequent periods of sleep and quiet rest [7, 9, 13].

**Fig 5.**
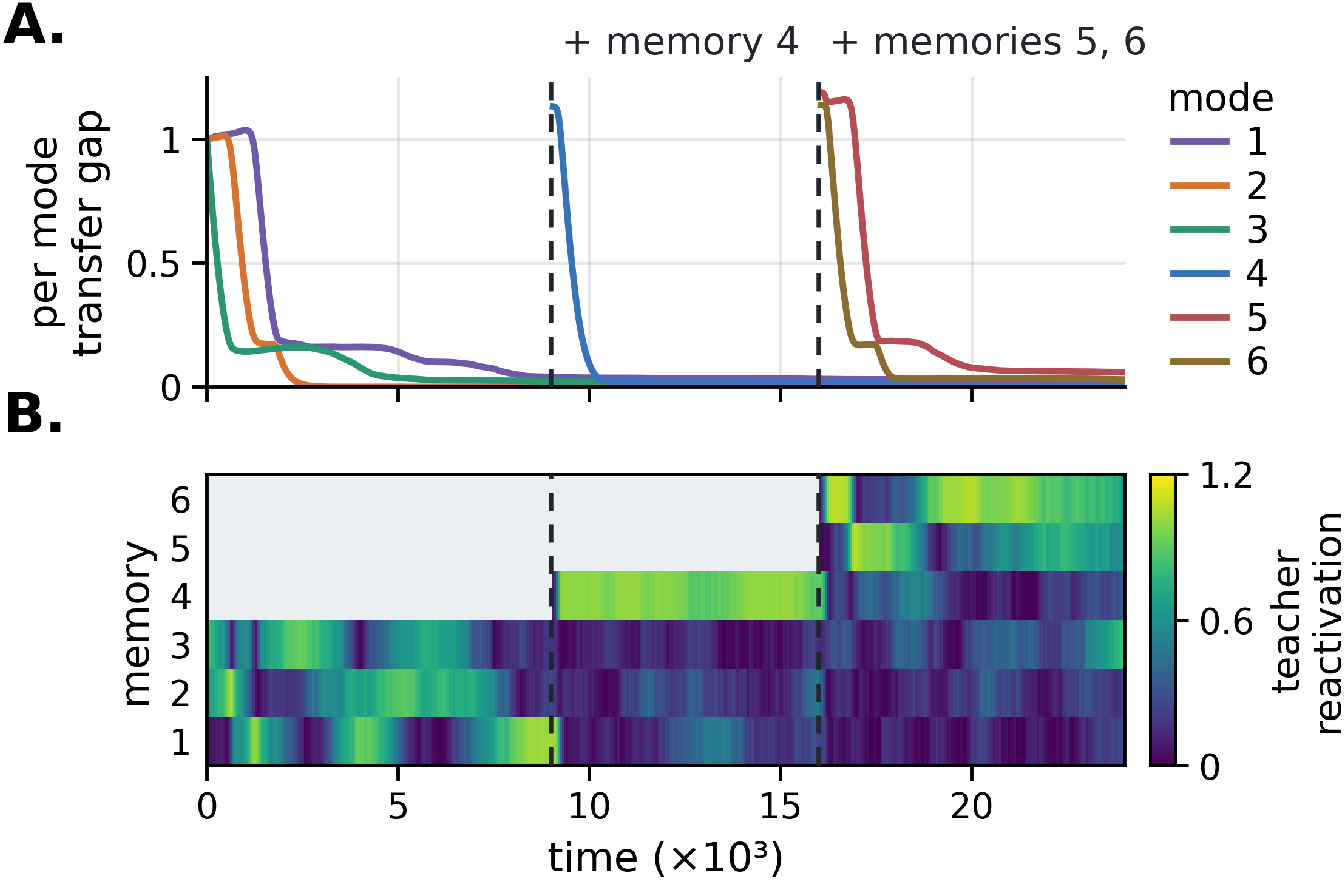
Incremental memory acquisition renews reactivation and memory transfer. A 50-unit teacher–student network was simulated in a single run while transferring 6 random memory patterns. The teacher initially stored *m*_1_, *m*_2_, and *m*_3_; it acquired *m*_4_ at *t* = 9000 and acquired *m*_5_ and *m*_6_ together at *t* = 16000 through interleaved teacher training on the whole corpus. Coupled intrinsic dynamics resumed after each acquisition without resetting student activity or recurrent weights (Materials and methods, Sections 5.3 and 5.4.4). Dashed lines mark the acquisition events. **A**, Per-mode transfer gaps during the simulation. Previously transferred modes keep a small error, whereas each newly supported mode introduces a large transfer gap that subsequently decreases. **B**, Memory coefficient magnitudes 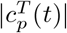, which measure the contribution of each stored pattern *m*^(*p*)^ to teacher activity *x*_*T*_ (*t*). Low values mean that the teacher does not reactivate the corresponding memory, whereas high values indicate strong reactivation. Newly added patterns dominate teacher activity immediately after their acquisition.

The observed dynamics are consistent with neocortex-dependent prioritisation [28]. In our model, the mismatch signal biases teacher activity toward teacher-supported modes that remain poorly represented by the student, and this bias weakens as those modes are learned. This provides a possible mechanistic account of the preferential reactivation of recently encoded hippocampal representations during post-learning sleep [5, 15], followed by reduced hippocampal involvement as memory retrieval becomes increasingly supported by neocortical circuits [10, 11]. Nevertheless, the model does not treat recency itself as the priority signal as sometimes proposed [25], nor does it require an explicit reactivation queue or novelty tag. Instead, it predicts that reactivation depends more directly on the mismatch between hippocampal support and cortical representation.

In summary, the coupled architecture transfers memory content from the teacher to the student during intrinsic activity. The student’s reconstruction errors on the teacher patterns decrease substantially but remain nonzero during the simulations, although the student already reconstructs recognisable versions of the stored patterns (Fig. 4). Tests involving classification of the reconstructed patterns or weaker query cues are still needed to determine whether, after transfer, the student can support the same computational role as the teacher in more challenging scenarios.

### 3.2 Fast transfer of correlated memory content

We next tested the transfer efficiency of our model on a memory corpus containing unequally sized groups of correlated patterns, both for a fixed corpus and when the teacher incrementally acquired new pattern groups.

To do so, we compared the following two protocols:

- **Intrinsic transfer (our model)**. A trained, frozen teacher storing the memory corpus was coupled to an initially empty student. Teacher and student activity evolved under the coupled intrinsic dynamics, and the student updated its recurrent weights after each integration step. In some experiments, memory patterns were added to the teacher memory corpus during transfer.
- **Uniform rehearsal**. At each plasticity event, one pattern was sampled uniformly from the memory corpus and clamped directly onto the student before a recurrent update was applied. In some experiments, memory patterns were added to the sampled corpus during training.

Uniform rehearsal emulates non-prioritised reactivation by sampling patterns uniformly. Comparing it with intrinsic transfer therefore tests whether mismatch-dependent dynamics improve transfer relative to uniform pattern sampling.

The results for the transfer of a correlated memory corpus are shown in Fig. 6.

**Fig 6.**
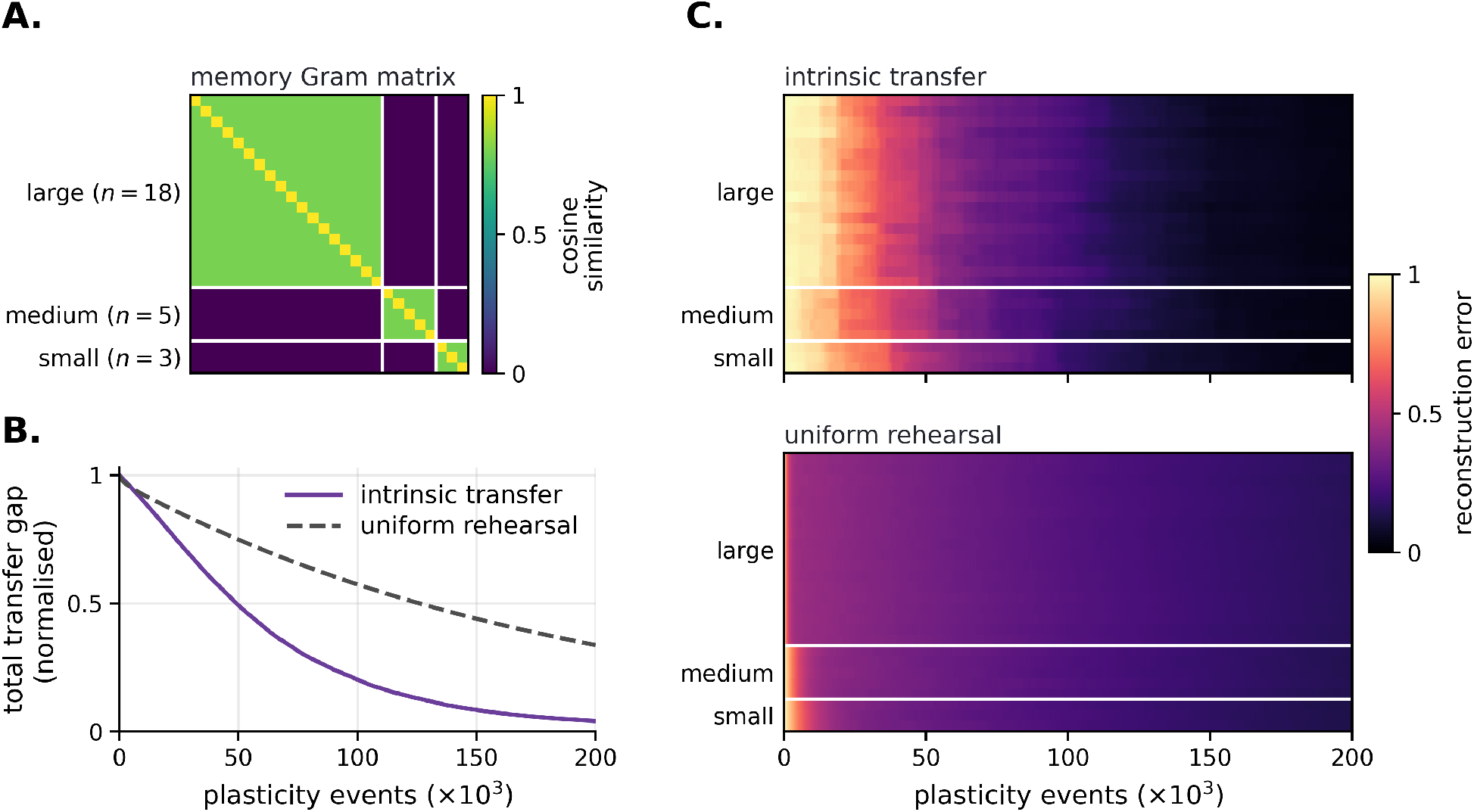
Intrinsic transfer accelerates transfer across unequally sized groups of correlated patterns. Both conditions used students initialised with *W*_*S*_ = 0, the same memory corpus, the same recurrent plasticity rule and plasticity rate. The corpus contained 26 patterns divided into three mutually orthogonal groups of 18, 5, and 3 patterns, with exact within-group cosine similarity *ρ* = 0.80. **A**, Memory Gram matrix. Each cell shows the cosine similarity between two patterns. **B**, Normalised total transfer gap 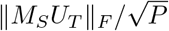 for intrinsic transfer (purple solid curve) and uniform rehearsal (grey dashed curve). Curves show the average over 20 simulations with different memory corpora. Intrinsic transfer reduced the total transfer gap faster than uniform rehearsal. **C**, Reconstruction errors for individual patterns during intrinsic transfer (top) and uniform rehearsal (bottom). Light colours indicate high reconstruction error and dark colours indicate accurate reconstruction. Intrinsic transfer reduced errors at similar rates across all three groups, whereas under uniform rehearsal, errors remained higher for longer in the smaller groups. Corpus construction, matched protocols, and measures are described in Materials and methods, Sections 5.5.1–5.5.3.

We considered a corpus of 26 patterns divided into three correlated groups of various sizes. We followed transfer at two levels. The individual-pattern residual used in the heatmaps asks whether the student can reconstruct each stored example. We also reported the normalised total transfer gap,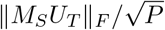. It gives equal weight to every independent memory-subspace mode and measures how much of the complete teacher memory subspace remains unsupported by the student.

The corpus construction, matched protocols and measures are described in Materials and methods, Sections 5.5.1, 5.5.2, and 5.5.3.

Intrinsic transfer produced a faster transfer than uniform rehearsal over the whole simulation (Fig. 6(B)). Fig. 6(C) shows that intrinsic transfer reduced reconstruction errors at similar rates across all three groups, despite their unequal sizes, whereas uniform rehearsal favoured learning in the largest group.

The difference in transfer rates reflects how each protocol selects the activity that drives learning. Uniform rehearsal samples individual patterns uniformly. Because patterns within each group are strongly correlated, these updates repeatedly reinforce their shared mode. This induces faster learning of over-represented groups. Intrinsic transfer does not depend on how often each pattern occurs in the corpus. As the student learns a teacher-supported mode, the mismatch for that mode contracts and its selection drive weakens, allowing activity to shift toward modes with larger remaining deficits. The result is consistent with deficit-dependent reactivation allocating plasticity according to reconstruction error across memory-subspace modes rather than the number of patterns in a correlated group.

Fig. 7 shows how within-group correlation affects the transfer rate of both protocols while group sizes remain fixed. Intrinsic transfer was unaffected by correlation: for every *ρ <* 1, the patterns of a group span the same subspace, so the teacher memory subspace and the intrinsic dynamics are unchanged. Only uniform rehearsal slowed as correlation increased. At low correlation, uniform rehearsal was faster, whereas increasing correlation shifted the advantage toward intrinsic transfer (Materials and methods, Sections 5.5.1–5.5.3).

**Fig 7.**
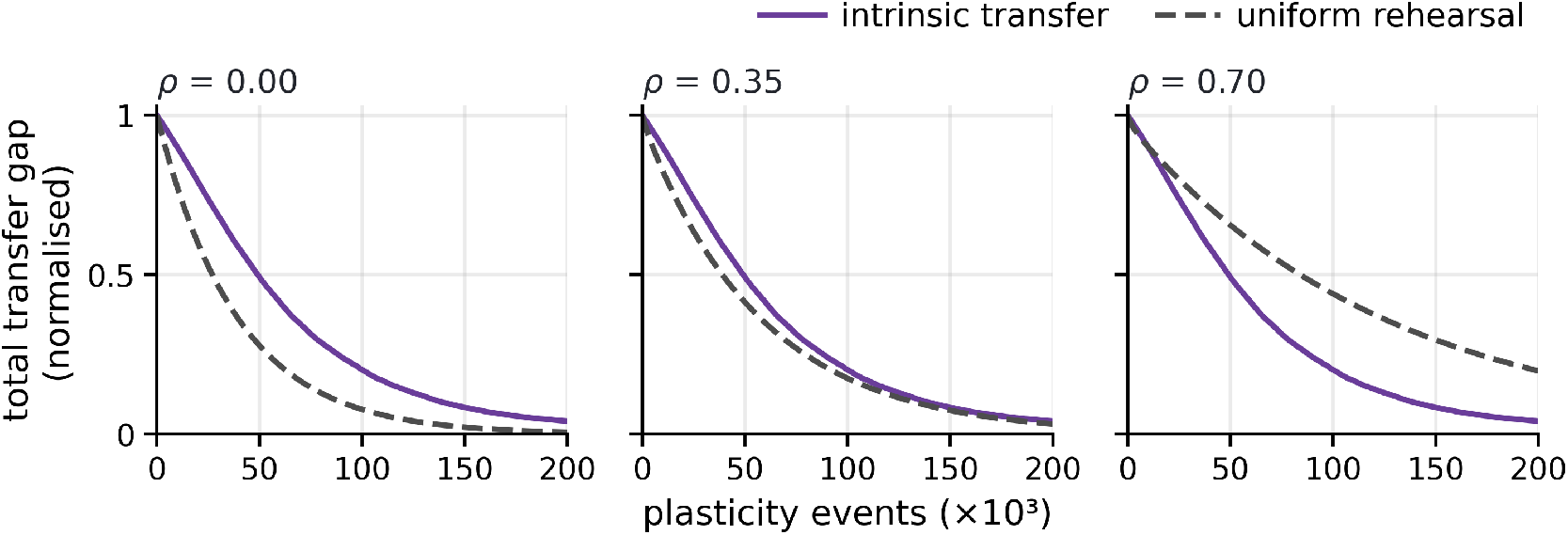
Increasing within-group correlation shifts the advantage from uniform rehearsal to intrinsic transfer. Same protocol as Fig. 6 for varying within-group cosine similarity *ρ* ∈ {0, 0.35, 0.70}. Each curve shows the normalised total transfer gap 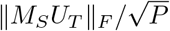, with lower values indicating more complete transfer. Grey dashed curves show transfer gap for uniform rehearsal and purple curves for intrinsic transfer. Curves show the average over 20 simulations. At *ρ* = 0, uniform rehearsal reduced the gap faster. At *ρ* = 0.35, the two protocols performed similarly. At *ρ* = 0.70, intrinsic transfer became faster. As correlation increased, uniform rehearsal slowed because uniformly sampled patterns produced increasingly overlapping updates, whereas the intrinsic-transfer curves remained unchanged. Corpus construction, matched protocols, and measures are described in Materials and methods, Sections 5.5.1–5.5.3.

Uniform rehearsal can be more efficient when correlations are weak because each update is based on an exact stored pattern. Intrinsic transfer updates the student while activity traverses the teacher memory subspace, so intermediate states can produce partly redundant updates. Uniform rehearsal therefore retains an advantage when uniform sampling already covers the memory subspace evenly.

We next asked whether this advantage persists when the teacher acquires new pattern groups during transfer (Fig. 8; Materials and methods, Sections 5.3 and 5.5.4). Each acquisition expanded the active teacher memory subspace and produced a new transfer gap. Intrinsic transfer reduced this gap faster than uniform rehearsal during every stage. These results are consistent with the mismatch-dependent dynamics redirecting activity and plasticity toward the newly added, unsupported content while preserving transfer of the groups already acquired by the student.

**Fig 8.**
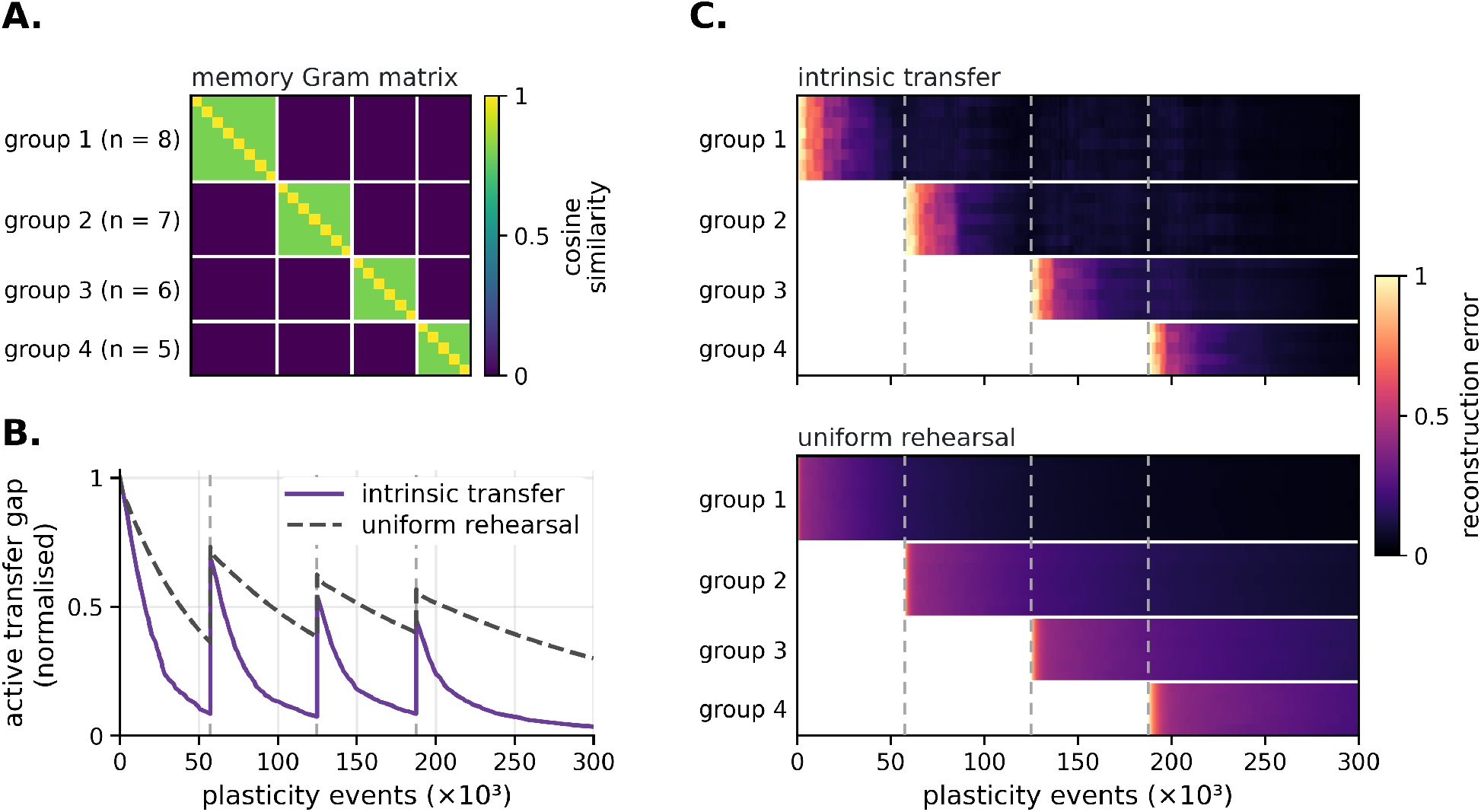
The intrinsic-transfer advantage persists during incremental acquisition of correlated pattern groups. As in the incremental-acquisition experiment in Fig. 5, patterns were added during a single transfer run by retraining the teacher on the enlarged corpus containing both the previously stored and newly introduced patterns. The students retained their activity and recurrent weights across acquisition events. The corpus contained 26 patterns arranged into four mutually orthogonal groups of 8, 7, 6, and 5 patterns, with exact within-group cosine similarity *ρ* = 0.80. Group 1 was present initially, and groups 2, 3, and 4 were introduced after 57,300, 124,800, and 187,600 plasticity events, respectively. Dashed vertical lines mark these acquisitions. **A**, Memory Gram matrix for the complete corpus. **B**, Normalised active-space transfer gap 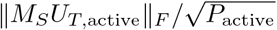 on the active patterns for intrinsic transfer (purple solid curve) and uniform rehearsal (grey dashed curve). Curves show the average over 20 simulations. Here, *U*_*T*,active_ is an orthonormal basis of the memory subspace supported by the teacher at that stage, and *P*_active_ is the number of patterns then available. Each acquisition expands the active memory subspace and produces an upward jump in the transfer gap. Intrinsic transfer reduces the gap faster during every stage. **C**, Reconstruction errors for individual patterns during intrinsic transfer (top) and uniform rehearsal (bottom). White areas mark patterns that have not yet been added to the active corpus. Once a group becomes active, its reconstruction errors decrease faster under intrinsic transfer, consistent with transfer being directed toward the newly unsupported content. The acquisition protocol and measures are described in Materials and methods, Sections 5.3, 5.5.3, and 5.5.4.

Together, these results show that mismatch-dependent intrinsic dynamics can transfer highly correlated memory content using fewer plasticity events than uniform rehearsal, both for a fixed corpus and when new patterns are incrementally added to the teacher. This advantage grows with correlation, because uniformly sampled patterns then produce increasingly overlapping updates. Such efficiency may matter in the brain, where neural activity carries a substantial metabolic cost and consolidation occurs during limited periods of sleep and quiet rest [1, 4, 5]. The same principle may help explain why reactivation favours memories that are new or weakly learned [5, 14]. Such memories may be less fully integrated into neocortical circuits [7], and prioritising them could accelerate their consolidation without repeatedly reinforcing traces that are already well established.

The next section provides a mathematical account of this qualitative behaviour. We ask whether the coupled dynamics can be understood as tracking the largest settled student free energy over the teacher memory subspace, thereby redirecting learning to where it is most needed.

### 3.3 Teacher activity tracks the largest settled student free energy, and plasticity reduces it

The preceding results showed that intrinsic transfer preferentially directs learning toward teacher-supported memory content that remains poorly stored by the student.

We next asked whether this prioritisation can be described in terms of the student’s free energy.

#### Hypothesis

During intrinsic activity, the teacher converges toward one of the states of its memory subspace for which settled student free energy is maximal. Student plasticity acting on this selected state then reduces its free energy.

This hypothesis is motivated by Theorem 1, presented below. Under simplifying assumptions, the theorem shows that the student’s influence drives teacher activity toward states with higher settled student free energy and may therefore bring activity toward a state that maximises it.

To obtain the following analytical result, we use the separation of timescales

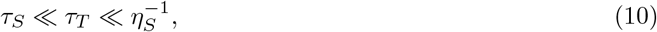

under which student activity relaxes faster than teacher activity, while both activity variables evolve faster than the student weights. This separation permits an adiabatic elimination of the student state for analysis. For fixed teacher activity *x*_*T*_ and student weights *W*_*S*_, let

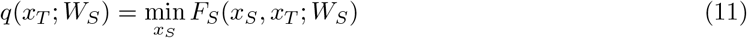

denote the student free energy remaining after its activity has settled. We call *q* the *settled student free energy*.

The teacher’s reachable memory sphere is

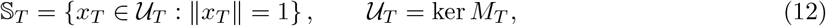

which consists of the network states that belong to the teacher memory subspace and lie on the activity sphere. As described in Section 2.3, teacher activity is normalised, and the parameters satisfy 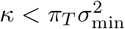 (Eq. (9)). The teacher’s recurrent correction therefore tends to keep activity near the teacher memory subspace. This choice makes the following on-subspace description a good approximation of the simulated dynamics.

The largest settled student free energy over the teacher memory subspace is

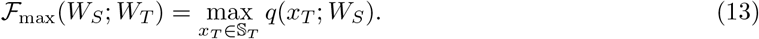

A high value of *ℱ*_max_ means that at least one state belonging to the teacher memory subspace induces a large student internal reconstruction error and therefore remains poorly stored by the student. Conversely, *ℱ*_max_ vanishes only when the student accurately stores every state in the teacher memory subspace. In this notation, the hypothesis is that teacher activity approaches a maximising state 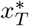 satisfying 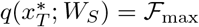.

We now present the analytical result that motivates the hypothesis.

**Theorem 1** (The settled interface drive points toward higher settled student free energy). *Let M*_*S*_ = *I* − *W*_*S*_ *and define*

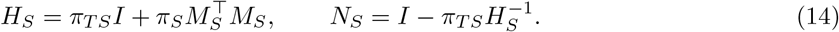

*For fixed x*_*T*_ *and W*_*S*_, *the student dynamics have the unique equilibrium*

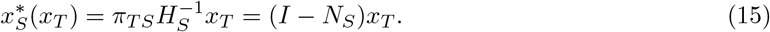

*In particular, the residual mismatch after the student has settled is*

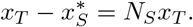

*At this equilibrium*,

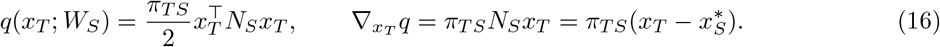

*Consequently, for κ >* 0, *the settled interface drive satisfies*

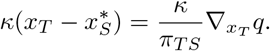

*It therefore points in the direction of increasing settled student free energy*.

*Proof*. Expanding the student free energy gives

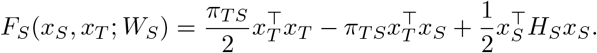

Its gradient and Hessian with respect to *x*_*S*_ are therefore

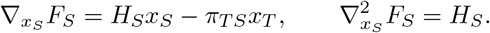

Since

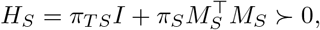

the free energy is strictly convex in *x*_*S*_, so the student dynamics have a unique equilibrium. Solving

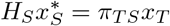

gives

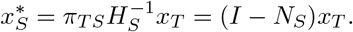

It follows directly that

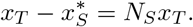

At equilibrium, 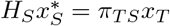. Hence

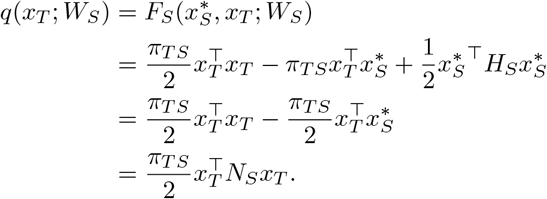

Because *H*_*S*_ is symmetric, *N*_*S*_ is also symmetric. Differentiating the last expression therefore yields

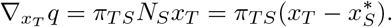

Finally,

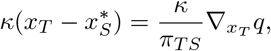

and *κ/π*_*T S*_ *>* 0, which proves the result.

This result shows that the interface-error drive acting on teacher activity is aligned with the direction of gradient ascent on the settled student free energy.

We then tested our hypothesis by tracking the angular distance between the teacher state and the nearest maximising state. Fig. 9 shows the results.

**Fig 9.**
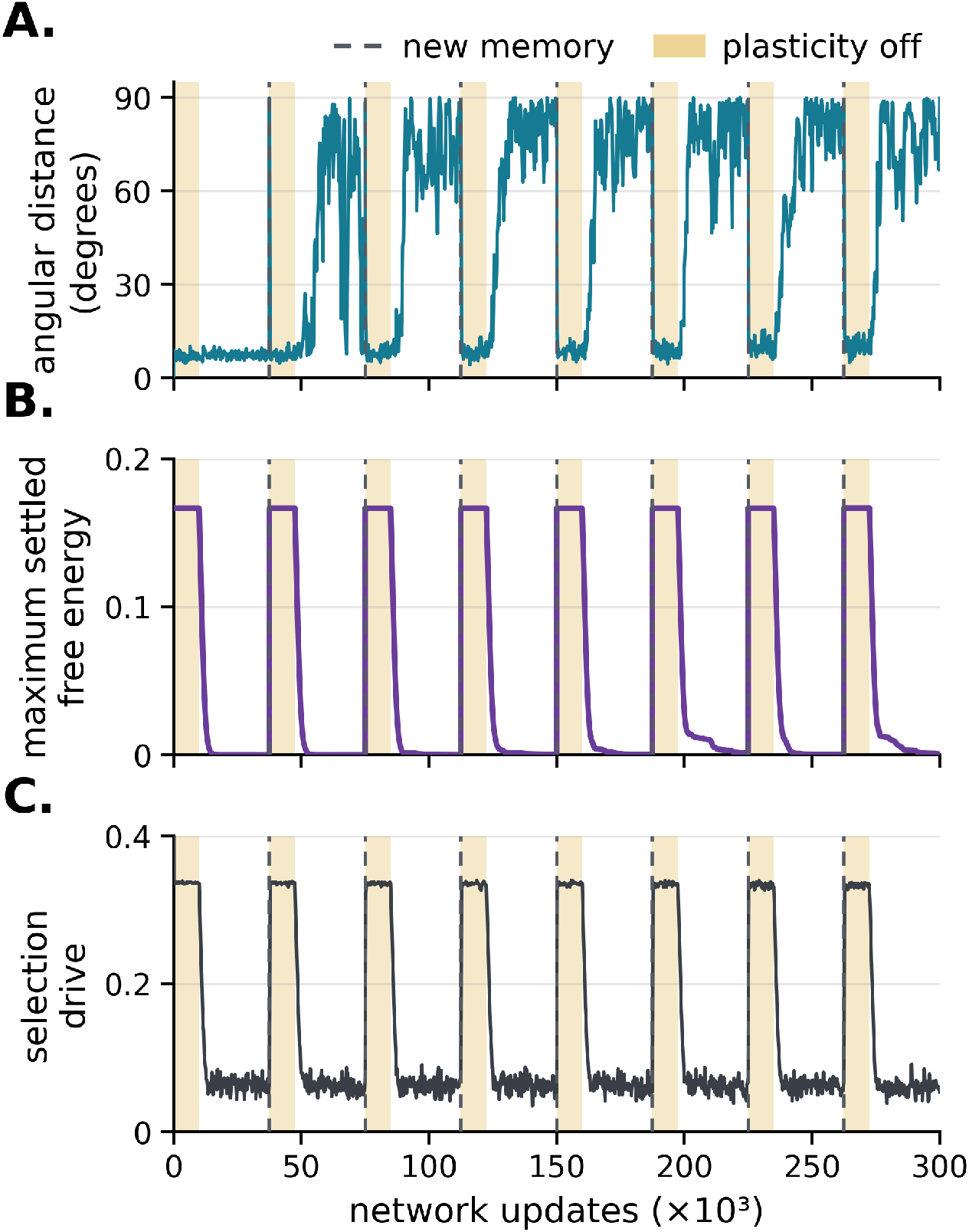
Intrinsic activity approaches the states that maximise settled student free energy, and student plasticity then reduces it. A 24-unit coupled teacher–student network incorporated eight orthogonal patterns incrementally during a single run. The teacher initially stored one pattern and acquired the remaining patterns individually. Teacher activity, student activity, and student weights were preserved across acquisitions. Dashed vertical lines mark acquisition events. Yellow intervals mark the first 10,000 events of each stage, during which student plasticity was disabled while coupled activity continued to evolve. The reference calculation and simulation protocol are described in Materials and methods, Sections 5.6.1 and 5.6.2. **A**, Angular distance between teacher activity and the nearest state maximising settled student free energy over the teacher memory subspace. Smaller angles indicate that teacher activity more closely tracks a state that maximises settled student free energy. After each acquisition, activity approached a maximising state during the frozen-weight interval. **B**, Largest settled student free energy over the teacher memory subspace. Each acquisition introduced a new unsupported memory mode and abruptly increased the maximum. The maximum remained constant while plasticity was disabled and decreased when student plasticity resumed. **C**, Teacher-side selection drive *κ* ∥*ε*_*T S*_∥ _2_, the magnitude of the interface-error drive acting on the teacher. The selection drive remained constant while plasticity was disabled and decreased when student plasticity resumed.

In these simulations, the teacher acquired eight mutually orthogonal patterns one at a time (Materials and methods, Section 5.3). We refer to the interval between two successive acquisitions as a stage; the first stage begins at the start of the run, with a single stored pattern. At the start of each stage, student plasticity was disabled for a fixed interval while the coupled activity dynamics continued to evolve. These frozen-weight intervals separate teacher-state selection by intrinsic activity from student learning (Materials and methods, Section 5.6.2). We call any state of the teacher memory subspace that maximises the settled student free energy a maximising state. Fig. 9(A) shows the angle between teacher activity and the nearest maximising state, (B) shows *ℱ* _max_, and (C) shows the teacher-side selection drive.

Each stage unfolded in three phases. First, the acquisition introduced a new unsupported memory mode, which abruptly increased both *ℱ*_max_ and the selection drive (Fig. 9(B, C)). Because the teacher state lay near the previous memory subspace, and the new mode was orthogonal to all previously stored patterns, each stage began with the teacher far from the maximising state, at an angle close to 90^*°*^. The first stage is an exception for which the one-dimensional memory subspace kept the angle small throughout.

Second, during the frozen-weight interval, *ℱ*_max_ and the selection drive remained constant, and the angle dropped rapidly below about 10^*°*^ as the teacher state converged toward the maximising state (Fig. 9(A)).

Third, once plasticity resumed, the student acquired the new memory mode: *ℱ*_max_ decreased toward zero and the selection drive fell to a small residual value. As *ℱ*_max_ approached zero, the angle increased again, fluctuating mostly between 60^*°*^ and 90^*°*^. This increase likely reflects two combined effects. With little selection drive remaining, noise-induced exploration moved the teacher state randomly within its memory subspace. In addition, all memory modes then had nearly equal, near-zero settled free energy, so the maximising state was no longer distinguished from other states of the memory subspace and was determined by small residual differences. The angle therefore ceases to be informative once the newly acquired pattern has been transferred to the student.

This cycle repeated at every acquisition: new teacher memory content increased the largest deficit and the selection drive, intrinsic activity aligned the teacher with the corresponding maximising state, and student plasticity then reduced its free energy.

These results support our hypothesis. Student relaxation converts the interface error into a drive pushing the teacher toward states with higher settled student free energy. Teacher activity follows this signal, and local student plasticity then reduces free energy through memory transfer.

## 4 Discussion

We introduced a network architecture grounded in predictive coding (PC) that enables autonomous memory transfer between neural memory systems. A teacher network, interpreted as a hippocampus-like structure, first stores a set of memory patterns during a phase analogous to wakefulness. During subsequent intrinsic activity, this memory content is transferred to a student network corresponding to the neocortex.

These dynamics arise from the asymmetric coupling between the two networks. Effective top-down inhibition from the student biases teacher activity toward regions of the teacher’s memory subspace that the student reconstructs poorly. The selected teacher activity then drives inference in the student, and local plasticity progressively encodes the corresponding memory content in the student’s recurrent weights.

The principal novelty of the proposed architecture is its use of interface-error feedback to prioritise memory reactivation according to the student’s remaining representational deficits. Local mismatch computations have been observed in cortical circuits [48, 49]. Assuming that similar circuits compute the hippocampal-neocortical interface error, the proposed architecture provides a circuit-level candidate mechanism for reactivation prioritisation and efficient offline memory content transfer.

The proposed mechanism accounts for several reported features of reactivation prioritisation. First, reactivation preferentially targets recently acquired memories over older ones. In the brain, hippocampal ensembles encoding recent experiences reactivate most strongly shortly after encoding, with reactivation declining over the following hours [3,15,50]. In the model, each newly stored hippocampal memory creates a mismatch between its hippocampal and neocortical representations, biasing reactivation toward that memory. As neocortical support develops, this mismatch diminishes and the memory is prioritised less, consistent with the observed time course of consolidation in the brain. Second, reactivation favours weakly learned memories. Schapiro et al. showed that, during awake rest after learning, the human hippocampus preferentially reactivated items that participants remembered less well, and that this reactivation predicted memory improvement over a subsequent interval [14]. In our framework, a weakly learned memory corresponds to one whose neocortical representation remains incomplete. It therefore induces a large interface error and continues to be prioritised for reactivation, whereas well-learned memories are progressively reactivated less. In both cases, priority in our model is not a property of the hippocampal trace alone, such as its recency or strength, but emerges from the relation between its hippocampal and neocortical representations.

Several computational accounts have proposed functional criteria for prioritising reactivation or consolidation, including prediction error, contribution to generalisation, memory reliability, and expected utility [21, 23, 24, 51]. Our architecture offers a complementary account that shows how a priority signal can arise from local dynamics inherent to the PC framework and subsequently bias neuronal activity.

The proposed architecture builds on the asymmetric interaction between the PFC and CA1 observed during sleep [26, 27]. A further implication of the model is that prefrontal control mechanisms engaged during wakefulness may be repurposed during sleep to regulate memory reactivation. During wakefulness, prefrontal control can suppress unwanted or competing hippocampal retrievals [52, 53]. This top-down control is potentially mediated by both direct and indirect pathways from the PFC to the hippocampus. Long-range GABAergic projections from the PFC to the hippocampus preferentially inhibit VIP-expressing interneurons, reducing local disinhibition and thereby increasing feedforward inhibition in CA1 [54]. Excitatory prefrontal projection neurons may also influence the hippocampus indirectly through the nucleus reuniens, whose hippocampal projections activate CA1 interneurons and generate polysynaptic inhibition [31, 32]. During wakefulness, behavioural goals determine which retrievals are suppressed, whereas our model suggests that, during sleep, suppression may instead depend on how well the retrieved content is already represented in the neocortex.

The model further predicts that cortical memory representations regulate hippocampal reactivation, a hypothesis that can be tested experimentally. Previous work has shown that PFC engrams undergo functional maturation during systems consolidation, while hippocampal engram cells become functionally silent over time [11, 55]. These findings are compatible with our model, but do not establish whether spontaneous hippocampal reactivation is regulated by the functional state of the corresponding cortical engram. Our model predicts such a content-specific relation: reversible silencing of a mature cortical engram during sleep, using activity-dependent tagging and optogenetic inhibition [56, 57], should selectively increase reactivation of hippocampal engram cells recruited during the same learning episode. Conversely, repeated optogenetic reactivation of a tagged cortical engram during post-learning sleep has been shown to accelerate systems consolidation [58]. Our model predicts that this accelerated cortical consolidation should subsequently reduce reactivation of hippocampal engram cells recruited during the same learning episode. These experiments would distinguish the proposed feedback mechanism from the time-dependent shift from hippocampal to neocortical memory dependence described by classical accounts of systems consolidation.

Our model rests on several simplifications. First, hippocampal and neocortical neurons are in one-to-one correspondence, so both networks represent stored patterns in the same neural code. Hippocampal indexing theory proposes instead that the hippocampus stores not a copy of neocortical content but an index of it [33, 34]. This index is a pattern of hippocampal activity linked to the neocortical neurons that were active during an experience, and reactivating it reinstates the corresponding neocortical pattern. Consistent with this view, silencing hippocampal neurons active during learning prevents the reinstatement of cortical neurons active at encoding and impairs retrieval [59]; see [60] for a review.

A more biologically faithful version of our model would give the hippocampus its own neural code, for example a sparse code in which similar experiences are represented by weakly overlapping activity patterns [45, 61]. The hippocampal and neocortical codes would then be linked by learned projections. Hippocampal-to-neocortical projections would map each index onto its neocortical pattern, and neocortical-to-hippocampal projections would carry the selective inhibitory feedback from the student to the teacher. These projections could be learned during encoding, as the index is formed, or during earlier periods of spontaneous activity.

Our model corresponds to the special case in which both projections reduce to a fixed one-to-one mapping. The hippocampus then drives the neocortex toward exactly the pattern it stores, and each student neuron inhibits only its counterpart teacher neuron. We restrict the model to this case because it greatly simplifies the mathematical analysis.

Anatomically, the hippocampus and the neocortex are linked by direct projections in both directions: CA1 and the subiculum project to the prefrontal cortex, and long-range GABAergic prefrontal neurons project to the hippocampus [30, 54]. The two systems also interact through polysynaptic routes, with the entorhinal cortex as the principal relay and the nucleus reuniens mediating indirect prefrontal influences on the hippocampus [29, 31, 32]. Our model abstracts away these relays to focus on the core dynamics of memory transfer.

Memory content may also be transformed during consolidation, notably during sleep. Memories can be integrated with existing knowledge, and regularities can be extracted across experiences, producing more schematic or gist-like representations [9,62–64]. Computational models suggest that these transformations support abstraction and generalisation [20, 23]. Our model does not include them: the student learns to reconstruct the teacher memory subspace, without transforming or abstracting its content.

Finally, the teacher and student networks are linear. Under this assumption, the teacher stores the entire linear span of its stored patterns, and transfer consists of making that same subspace reconstructable by the student. This setting isolates the proposed prioritisation mechanism but omits nonlinear neuronal responses, spiking activity, and the distinct architectures of the two systems. The model also does not account for factors known to modulate the prioritisation of reactivation, such as reward [65, 66].

Future work should therefore investigate whether transfer-gap-dependent prioritisation persists in networks with nonlinear activation functions or when the teacher and student use different neural codes and must learn a mapping between them. Future work should also examine how behavioural relevance and neuromodulatory signals could interact with our proposed mechanism in controlling reactivation.

Together, our results identify neocortical feedback as a possible mechanism by which hippocampal reactivation is progressively redirected away from memory content already supported by the neocortex and toward content whose neocortical representation remains incomplete. More generally, this mechanism frames reactivation prioritisation as a transfer of memory content across memory systems, in which reactivation priority reflects the extent to which hippocampal memory content has been incorporated into neocortical representations.

## 5 Materials and methods

### 5.1 Numerical simulation and network initialisation

Teacher and student networks have the same number *d* of neurons. Unless stated otherwise, the student starts with *W*_*S*_ = 0 and *x*_*S*_ = 0. At the start of each independent simulation, we draw the teacher state isotropically within the active teacher memory subspace and normalise it to unit length. The diagonal of every recurrent weight matrix remains fixed at zero throughout, so neither network contains autapses. Table 1 summarises the parameters and numerical conventions that remain fixed across simulations throughout the Results section.

**Table 1.** Parameters and numerical conventions held fixed across intrinsic-transfer simulations.

| Quantity | Fixed choice |
| --- | --- |
| Activation function | $f(x) = x$ |
| Student recurrent precision | $\pi_S = 0.5$ |
| Teacher–student interface precision | $\pi_{TS} = 1$ |
| Initial teacher activity | Normalised isotropic Gaussian draw in the active teacher memory subspace |
| Teacher noise amplitude | $\sigma_\xi = 0.05$ |
| Interface coupling | $\kappa = 0.33 \pi_T \sigma_{\min}^2$ (Eq. (9)) |
| Recurrent self-connections | $(W_T)_{ii} = (W_S)_{ii} = 0$ |
| Numerical precision | 64-bit floating point |

We integrate the deterministic activity dynamics and student plasticity by the explicit Euler method with time step Δ*t*. We integrate the stochastic teacher term by Euler–Maruyama: at every step, an independent vector *z*_*n*_ ~ *N* (0, *I*_*d*_) contributes 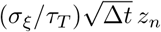 to teacher activity. After this increment, we rescale teacher activity to unit norm.

The student weights are updated as

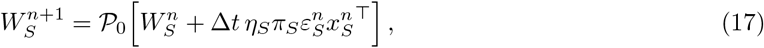

where *P* _0_(*A*) denotes the projection that sets the diagonal entries of *A* to zero. The simulations use 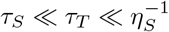 so that student activity responds faster than teacher activity and both activity dynamics are faster than synaptic plasticity.

The integration step, time constants, and plasticity rates depend on the experimental protocol and are therefore reported separately.

### 5.2 Interleaved training of the teacher

Before allowing the memory transfer dynamics to occur, we train the teacher independently, without any coupling to a student network.

The training corpus is *ℳ* = [*m*^(1)^, …, *m*^(*P*)^] ∈ ℝ^*d×P*^, where each column is a unit-normalised memory pattern. Initial teacher training begins from *W*_*T*_ = 0. During each training epoch, we randomly permute the pattern order and successively clamp teacher activity to every pattern in the corpus. For a clamped pattern *m*^(*p*)^, the recurrent error is *ε*_*T*_ = (*I* − *W*_*T*_)*m*^(*p*)^, and we apply one local plasticity step per clamped presentation,

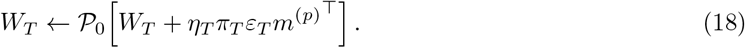

We repeat interleaving until the largest pattern residual is at most *δ*_*T*_, subject to a maximum of *N*_ep_ epochs. We do not update the student activity or weights during this phase. Algorithm 1 summarises the procedure, and Table 2 displays its parameters and fixed conventions.

**Table 2.**
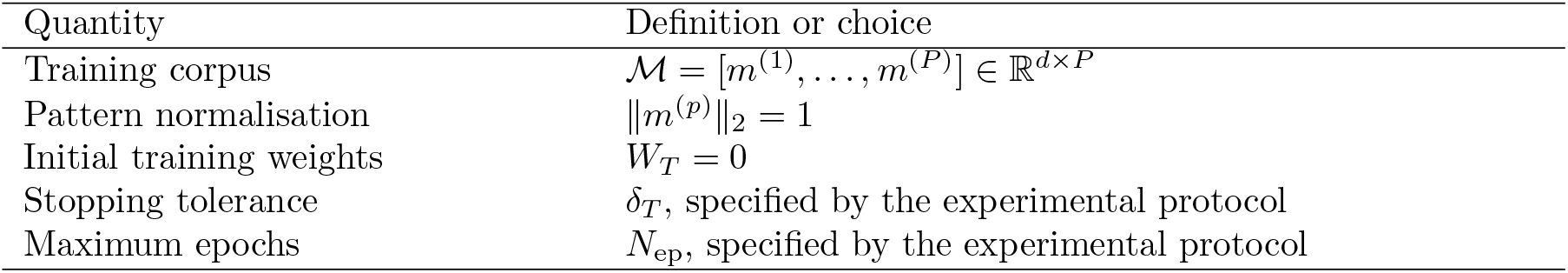
Parameters and conventions for interleaved teacher training.

#### Algorithm 1

Interleaved training of the teacher network

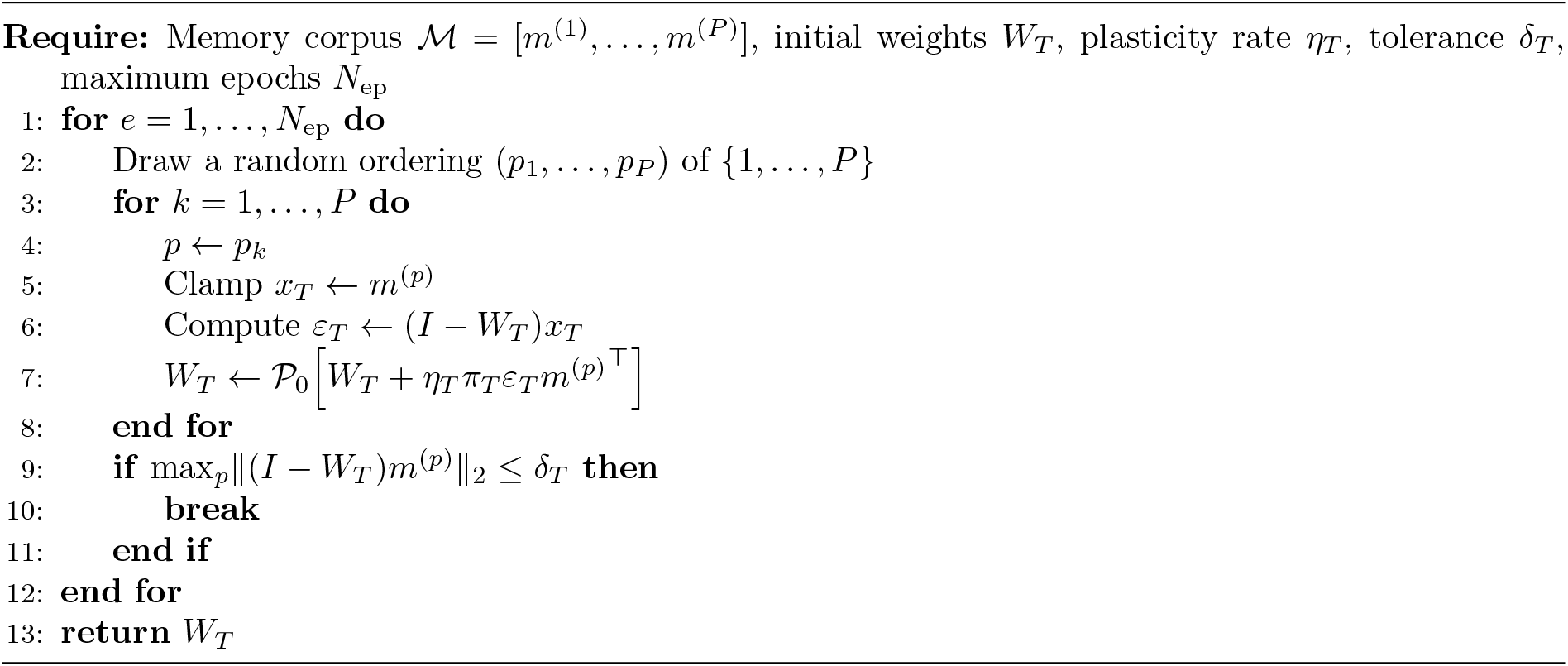

### 5.3 Incremental memory acquisition during transfer

For experiments in which the teacher acquires new patterns during transfer, we pause intrinsic simulation at each acquisition time. We append the new patterns to the active teacher memory corpus and retrain the teacher by Algorithm 1 on the complete enlarged corpus, including both previous and newly added patterns. Retraining begins from the current *W*_*T*_ rather than from zero. The interface and student plasticity remain inactive during this phase. We preserve *W*_*S*_ and both activity states across the acquisition event. After retraining, intrinsic activity resumes from the state reached immediately before the pause, with the new teacher weights and otherwise unchanged simulation parameters. The acquisition event therefore changes the teacher memory subspace without resetting the student or restarting the transfer trajectory.

### 5.4 Reactivation and memory content transfer

#### 5.4.1 MNIST transfer protocol

We prepared the MNIST memory corpus as specified in Table 3, vectorising and normalising each selected image to unit Euclidean norm. We trained the teacher by Algorithm 1 and then ran memory transfer through intrinsic coupled dynamics. Table 3 gives the experiment-specific parameters; Table 1 lists the remaining fixed conventions.

**Table 3.** Parameters for the MNIST transfer experiment.

| Quantity | Value |
| --- | --- |
| MNIST samples | one sample for each label in $\{0, 2, 7\}$ |
| Image and network dimension | $28 \times 28$ pixels; $d = 784$ |
| Teacher-training tolerance | $\delta_T = 10^{-8}$ |
| Maximum teacher-training epochs | $N_{\text{ep}} = 3000$ |
| Teacher recurrent precision | $\pi_T = 1$ |
| Teacher and student time constants | $\tau_T = 10, \tau_S = 1$ |
| Plasticity rate | $\eta_S = \eta_T = 0.005$ |
| Integration step | $\Delta t = 0.5$ |
| Simulation duration | 7,500 time units |

#### 5.4.2 Transfer and reconstruction measures

Let *ℳ* = [*m*^(1)^, …, *m*^(*P*)^] denote the teacher memory corpus, with one pattern per column. Let *r* = rank(*ℳ*) and *U*_*T*_ = [*u*_1_, …, *u*_*r*_] ∈ ℝ^*d×r*^. Each column *u*_*j*_ is a unit vector, distinct columns are orthogonal, and together the columns span the same space as the stored patterns: 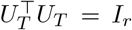 and span(*U*_*T*_) = span(*ℳ*).

At each recorded state, we calculate the teacher and student memory coefficients and the restricted student error matrix as

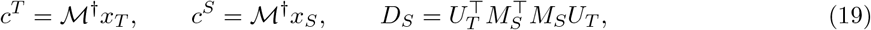

where *ℳ*^*†*^ is the Moore–Penrose pseudoinverse. The reactivation plots show 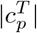 and the magnitude gap 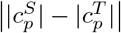. Per-mode transfer gaps are the eigenvalues of *D*_*S*_, and the total gap is ∥*M*_*S*_*U*_*T*_ ∥_*F*_. We also record the magnitude of student weight updates 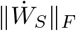 and teacher-side selection drive *κ*∥*ε*_*T S*_∥_2_. For each pattern, the direct reconstruction residual is *R*_*p*_(*t*) = ∥*m*^(*p*)^ − *W*_*S*_(*t*)*m*^(*p*)^∥_2_*/*∥*m*^(*p*)^∥_2_.

#### 5.4.3 Clamped reconstruction probes

We probed the isolated teacher and student networks under the conditions in Table 4. For each probe, we clamped the visible neurons to the corresponding part of the pattern, allowed the remaining activity to evolve until convergence, and recorded the final activity of all neurons.

**Table 4.** Parameters for the reconstruction probes.

| Quantity | Value |
| --- | --- |
| Student-weight checkpoints | $t \in \{1550, 6500\}$ |
| Visible neurons | Upper 14 image rows |
| Hidden-unit initialisation | $x_{\alpha, H} = 0$ |

#### 5.4.4 Incremental-acquisition protocol

For Fig. 5, we generated and normalised the memory corpus as specified in Table 5. Each acquisition followed Section 5.3, with the schedule and parameters given in the same table. We calculated per-mode gaps from *D*_*S*_ restricted to the active teacher memory subspace.

**Table 5.** Parameters for the incremental-acquisition experiment.

| Quantity | Value |
| --- | --- |
| Memory corpus | $P = 6$ draws from $\mathcal{N}(0, I_{50})$, each normalised to unit length |
| Initial teacher corpus | $\{m_1, m_2, m_3\}$ |
| First acquisition | $m_4$ at $t = 9000$ |
| Second acquisition | $m_5, m_6$ at $t = 16000$ |
| Teacher-retraining rate | $\eta_T = 0.30$ |
| Teacher-retraining tolerance | $\delta_T = 10^{-8}$ |
| Maximum retraining epochs | $N_{\text{ep}} = 3000$ |
| Simulation duration | 24,000 time units |
| Other simulation parameters | Table 3 |

**Table 6.**
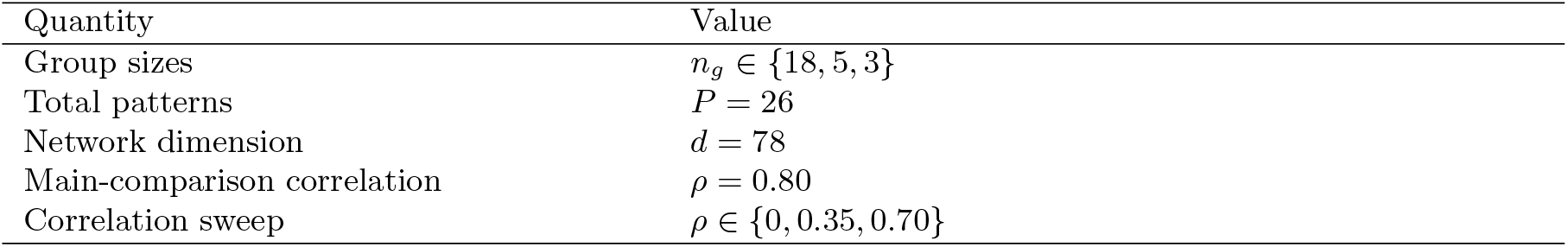
Parameters for the correlated-pattern corpus.

| Quantity | Value |
| --- | --- |
| Group sizes | $n_g \in \{18, 5, 3\}$ |
| Total patterns | $P = 26$ |
| Network dimension | $d = 78$ |
| Main-comparison correlation | $\rho = 0.80$ |
| Correlation sweep | $\rho \in \{0, 0.35, 0.70\}$ |

### 5.5 Fast transfer of correlated memory content

#### 5.5.1 Generating a memory corpus

For each group *g*, let *n*_*g*_ denote the number of memory patterns in that group. We assign each group to a separate *n*_*g*_-dimensional subspace. The columns of 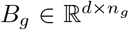 form an orthonormal basis for this subspace, and the bases assigned to different groups are mutually orthogonal. Within group *g*, we set the target Gram matrix and the corresponding memory matrix to

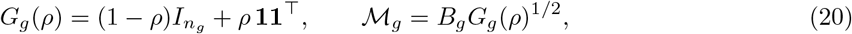

where *G*_*g*_(*ρ*)^1*/*2^ is the positive symmetric square root. The diagonal entries of *G*_*g*_ are one and its off-diagonal entries are *ρ*. Because 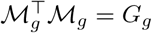, each column of *ℳ*_*g*_ has unit norm, and any two patterns in the same group have cosine similarity *ρ*. Patterns from different groups are orthogonal. We form the complete corpus as *ℳ* = [*ℳ*_1_, …, *ℳ*_*G*_].

#### 5.5.2 Matched transfer protocols

##### Uniform rehearsal

We trained the student directly on the complete corpus, without a teacher network or teacher–student interface. Each plasticity event independently sampled one pattern *m*^(*p*)^ with uniform probability, clamped it as the student activity, and applied one plasticity step of the same form as Eq. (18),

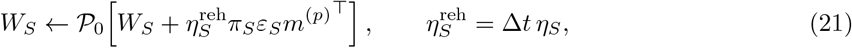

so that each rehearsal event applies the same weight step as one intrinsic-transfer event. Table 7 gives the protocol parameters; the fixed student conventions follow Table 1.

**Table 7.**
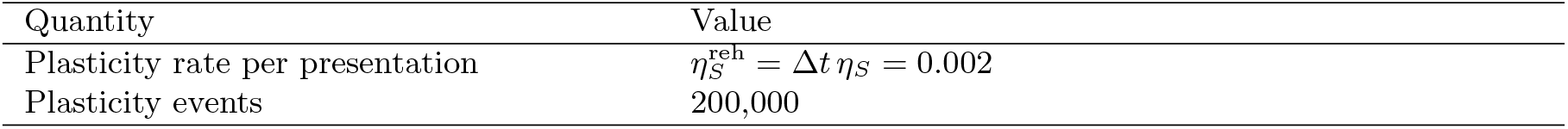
Parameters for uniform rehearsal of the correlated-pattern corpus.

| Quantity | Value |
| --- | --- |
| Plasticity rate per presentation | $\eta_S^{\text{reh}} = \Delta t \eta_S = 0.002$ |
| Plasticity events | 200,000 |

##### Intrinsic transfer

We first trained the teacher on the complete corpus using Algorithm 1, froze *W*_*T*_, and simulated the coupled teacher–student network. Each plasticity event consisted of one integration step of the full coupled dynamics, which simulated teacher activity, student activity, and the student recurrent weights. Table 8 gives the protocol parameters; the remaining conventions follow Table 1.

**Table 8.** Parameters for intrinsic transfer of the correlated-pattern corpus.

| Quantity | Value |
| --- | --- |
| Teacher training | $\eta_T = 0.005$, $\delta_T = 10^{-8}$, $N_{\text{ep}} = 5000$ |
| Teacher recurrent precision | $\pi_T = 1$ |
| Teacher and student time constants | $\tau_T = 1$, $\tau_S = 0.1$ |
| Student plasticity rate | $\eta_S = 0.04$ |
| Integration step | $\Delta t = 0.05$ |
| Plasticity events | 200,000 |

For intrinsic transfer, one plasticity event comprises one integration step of duration Δ*t* and one student-weight update; for uniform rehearsal, it comprises one clamped presentation and one student-weight update. Both protocols started from an empty student, and equal numbers of events correspond to equal numbers of student-weight updates of the same step size.

#### 5.5.3 Transfer measures

The individual-pattern heatmaps report the reconstruction residual *R*_*p*_(*t*) defined in Section 5.4.2. Using *M*_*S*_(*t*) = *I− W*_*S*_(*t*) and the teacher-space basis *U*_*T*_ defined in the same section, we calculate the normalised total transfer gap

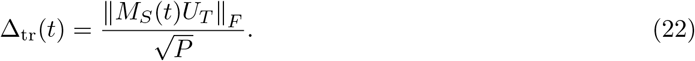

Because these corpora have *P* independent modes, Δ_tr_ gives equal weight to each mode.

For the transfer-gap curves in Figs. 6(B), 7, and 8(B), we averaged 20 simulations at each plotted time point. Standard-deviation bands are omitted because they are too small to resolve at the figure scale.

#### 5.5.4 Incremental acquisition of correlated groups

For Fig. 8, the teacher initially stored the first correlated group and acquired each remaining group according to Table 9. Teacher acquisition followed Section 5.3: we paused intrinsic simulation, retrained the teacher on the complete active corpus, and resumed without resetting activity or student weights. At the same events, the uniform-rehearsal corpus expanded to include the new group. Sampling remained uniform over the active patterns.

**Table 9.**
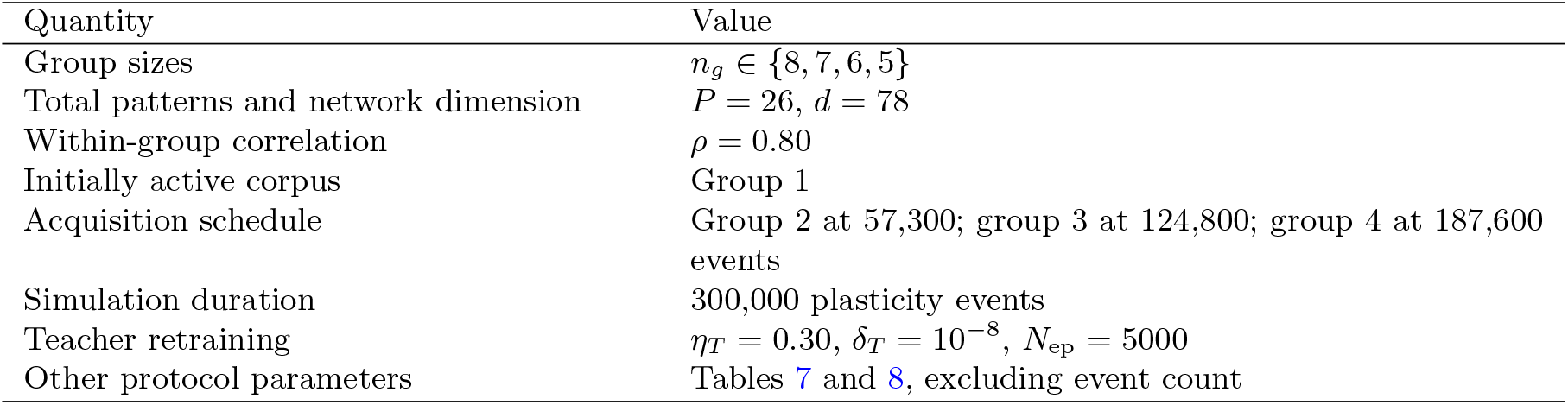
Parameters for incremental acquisition of correlated pattern groups.

| Quantity | Value |
| --- | --- |
| Group sizes | $n_g \in \{8, 7, 6, 5\}$ |
| Total patterns and network dimension | $P = 26, d = 78$ |
| Within-group correlation | $\rho = 0.80$ |
| Initially active corpus | Group 1 |
| Acquisition schedule | Group 2 at 57,300; group 3 at 124,800; group 4 at 187,600 events |
| Simulation duration | 300,000 plasticity events |
| Teacher retraining | $\eta_T = 0.30, \delta_T = 10^{-8}, N_{\text{ep}} = 5000$ |
| Other protocol parameters | Tables 7 and 8, excluding event count |

At each stage, let *U*_*T*,active_ span the active teacher memory subspace and let *P*_active_ be the number of active patterns. We measure

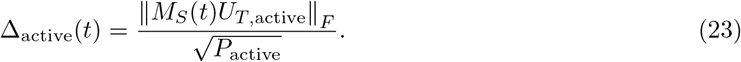

The individual-pattern heatmaps contain residuals only for patterns available at the corresponding stage.

### 5.6 Maximum-free-energy tracking during incremental acquisition

#### 5.6.1 Maximum-free-energy reference and tracking angle

At each measurement, let the columns of *U*_*T*_ form an orthonormal basis of the active teacher memory subspace. Using the operator *N*_*S*_ from Eq. (14), we restrict the settled student free energy to this space through

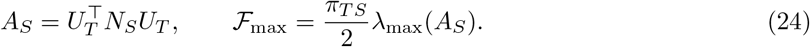

The leading eigenspace of *A*_*S*_ contains the coefficients of all teacher states that attain ℱ_max_. Let the columns of *Q*_*\**_ form an orthonormal basis of this eigenspace. We measure the angular distance between the simulated teacher activity and the nearest maximising state as

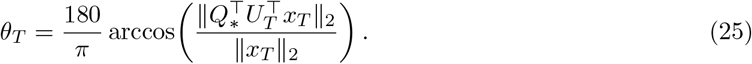

The full teacher activity enters this measure, so departures from the active memory subspace contribute to the angle. Using `numpy.isclose` (NumPy v1.26.0) [67], eigenvalues within relative tolerance 10^*−*7^ and absolute tolerance 10^*−*10^ of the largest eigenvalue are treated as part of the same maximising eigenspace.

#### 5.6.2 Incremental acquisition and plasticity-hold protocol

We generated eight mutually orthogonal, unit-normalised patterns for a 24-unit network. The teacher initially stored the first pattern and acquired the remaining patterns individually during a single intrinsic-transfer run. Each acquisition followed Section 5.3: we paused the simulation, retrained the teacher on the complete active corpus, and resumed without resetting teacher activity, student activity, or student weights.

At the beginning of every stage, including the initial stage, we set the student plasticity rate to zero for the fixed-weight interval given in Table 10. Teacher and student activity continued to follow the coupled dynamics during this interval. We recorded the teacher-side selection drive *κ* ∥ *ε*_*T S*_∥ _2_ as in Section 5.4.2. The remaining fixed conventions follow Table 1.

**Table 10.** Parameters for free-energy tracking during incremental acquisition.

| Quantity | Value |
| --- | --- |
| Memory corpus and network dimension | $P = 8$ orthogonal patterns, $d = 24$ |
| Acquisition schedule | One pattern every 37,500 events |
| Simulation duration | 300,000 plasticity events |
| Fixed-weight interval | First 10,000 events of each stage |
| Teacher retraining | $\eta_T = 0.075, \delta_T = 10^{-8}, N_{\text{ep}} = 5000$ |
| Coupled dynamics | $\pi_T = 2, \tau_T = 1, \tau_S = 0.1$ |
| Plasticity and integration step | $\eta_S = 0.04, \Delta t = 0.05$ |

